# Systemic effects of oral tolerance improve the healing of several and concomitant wounds in different body parts

**DOI:** 10.1101/2024.10.21.619410

**Authors:** Isabela Beatriz Cabacinha Nobrega, Angélica Vitória Souza Andrade, Thomson Junior Nyetem Bikat, Gustavo Motta Quintão, Geraldo Magela Azevedo Junior, Karen Franco Valência, Raquel Alves Costa, Claudia Rocha Carvalho

## Abstract

Throughout our lives, we are continually subjected to different situations that can result in several and concomitant wounds to different parts of our body. The healing of these wounds is essential to maintaining health. Inflammation is an important step in wound healing, but in cases of intense or prolonged inflammation, pathological scarring or non-closure of the wound may occur. Assuming that leukocytes participate in wound healing and that it is possible to intervene systemically with inflammation, we investigated a way to promote better repair of multiple wounds that may occur at the same time. Oral tolerance is an immunological phenomenon that result from protein intake and that have systemic effects on inflammation. Previous works have shown that parenteral injection of tolerated proteins reduces the inflammatory infiltrate and improves skin wound healing. Herein we tested whether the injection of tolerated proteins improves the healing of several wounds in different body parts, such as on the skin of the back and in the external ear (the auricle). To induce oral tolerance to ovalbumin (OVA), eight weeks old C57BL/6 mice drank egg white diluted 1:5 in water for 3 consecutive days. Control mice drank water. Seven days after oral treatment, mice were submitted to excisional injuries in the skin of the back (6 mm) and in the ears (4 mm). Minutes before the injuries, the mice received an intraperitoneal injection of OVA + Al(OH)_3_. Seven and 40 days after injuries, tissue samples were collected and processed for histological analysis of the wounds. The results show that the injection of OVA in animals that drank OVA reduced the inflammatory infiltrate in all lesions. Besides, injection of OVA in animals that drank OVA promoted better organization of the extracellular matrix, with thicker and intertwined collagen fibers in the neodermis, resulting in smaller scars in the skin. Furthermore, the healing area of the ears of OVA-tolerant animals showed chondrocyte aggregates and less obvious fibrous scar tissue compared with control animals. In conclusion, systemic effects of oral tolerance positively influenced the healing of several lesions in different body parts.

## Introduction

Multiple injuries can occur in different situations and may result from physical impacts, chemical products or pathological agents ^1,2^. In any situation, injuries result in disruption of the integrity of tissues and organs and, if not repaired in a timely manner, can have serious consequences for the body, including death. In humans, injuries are normally repaired by processes that replace the original tissue with a disordered extracellular matrix resulting in scars ^3,4^. On the skin, scars are easily visible as the closure of the lesions is normally not regenerative; dermal integrity is reestablished, but with replacement by scar tissue, a tissue morphologically and physiologically different from intact skin ^5^. In internal organs, scars can be noticed due to changes in their physiology.

Multiple injuries, resulting from accidents or situations caused by our way of life or conflict situations on our society, are frequent occurrences and generate great demands for health services ^1^. Fortunately, there are many ways to treat wounds and save lives. However, the occurrence of scars or delays in closing injuries are still common. Therefore, studies are still needed to generate effective treatments for closing wounds and reducing scars.

Inflammation is important for the initiation of repair and resolution of inflammation is necessary for wound closure. Prolonged inflammation can result in chronic wounds or hypertrophic scars ^5,6^. Once changes in the inflammatory phase of repair influence their subsequent phases, altering the inflammatory microenvironment with anti-inflammatory products and biomaterials is becoming an attractive approach in regenerative medicine ^7^.

It has long been known that one of the consequences of protein intake is oral tolerance, defined as refractoriness to subsequent immunization with the same protein ^8,9^. It is worth mentioning that, parenteral injection of tolerated proteins has systemic effects that inhibit inflammation and immunization to other agents injected simultaneously, 3 days before or shortly after ^10,11^. Injection of ovalbumin (OVA) or zein plus adjuvant, in animals fed with these proteins, reduces the inflammatory infiltrate in lesions caused by different agents, whether physical (punch injury) ^12,13^, chemical (produced by carragenin) ^14^ or biological agents (*Schistosoma manson*i eggs) ^15,16^. In cases of physical injuries, the positive effects of injecting tolerated proteins have been demonstrated in skin injuries and bone injuries, resulting in improved wound healing ^17^.

In the present work, the objective was to verify whether the injection of a tolerated protein improves the healing of several injuries, made at the same time and in different parts of the body. Following ethical principles, injuries were made with biopsy punch on the skin of the back and in the external ears of OVA-tolerant mice minutes after injecting OVA intraperitoneally (i.p.), and the wounds were analyzed 7 and 40 days after injuries.

## Material and methods

### Animals

Male C57BL/6 mice, eight weeks old, were obtained from Universidade Federal de Minas Gerais (UFMG), Brazil, and treated in accordance with the guidelines of the Institutional Animal Care and Use Committee. The animals were divided into groups of 5 animals each, for each time analyzed (N=5) and kept in a temperature-controlled environment at 24ºC, with a light cycle of 12 hours light and 12 hours dark.

### Treatment for induction of oral tolerance to ovalbumin (OVA)

The experimental group received as the only source of liquid, for 3 consecutive days, a solution of egg white diluted 1:5 in water (Figure 1a). The egg white solution was changed every day in the early evening. The control groups drank tap water.

**Figure 1.**
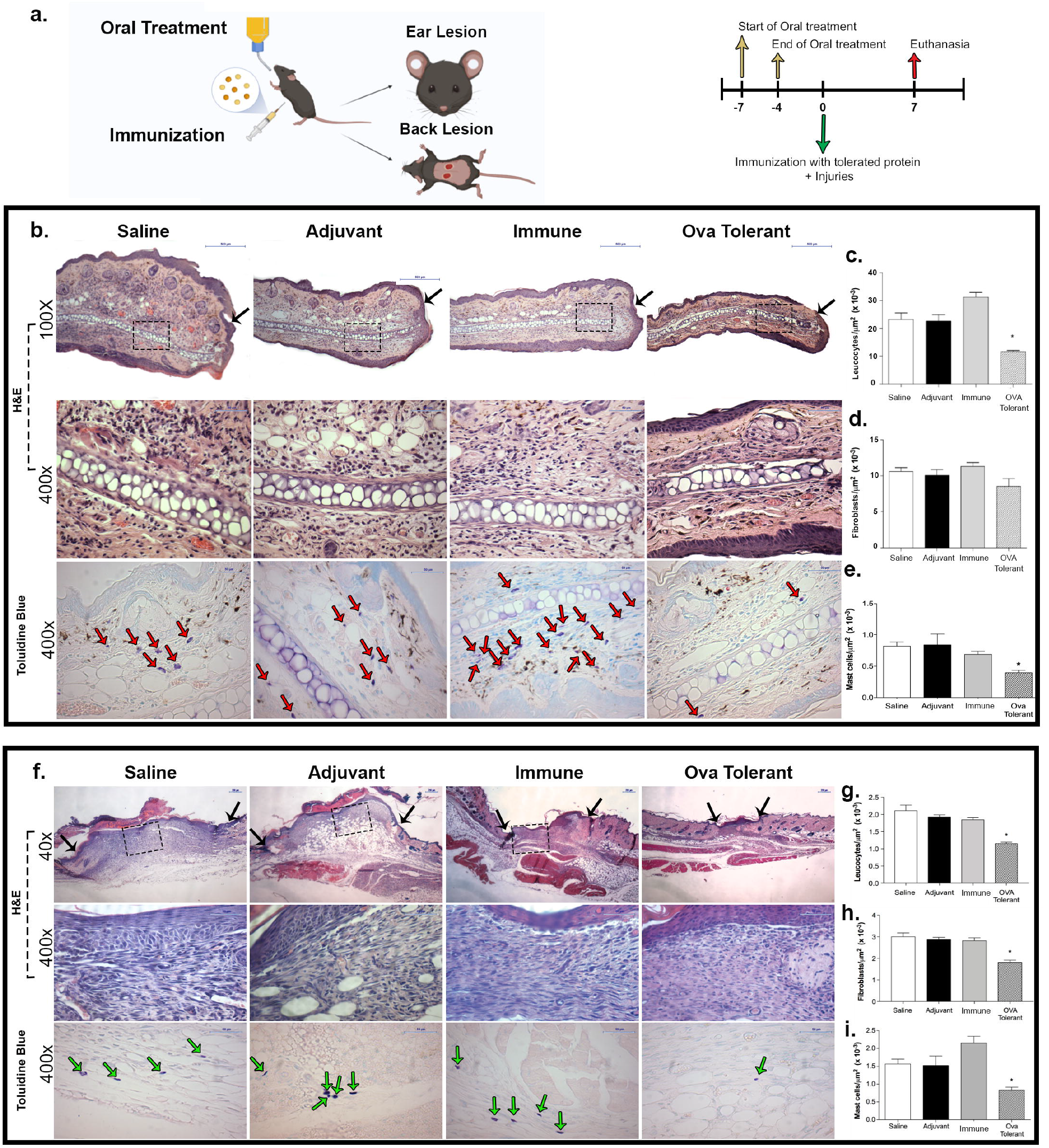
The i.p injection of OVA in animals fed with this protein reduced the inflammatory infiltrate in the wound beds of the ears and back skin. (a) Experimental design for analysis at day 7; (b) Photomicrographs of ear sections, 7 days post-injury, stained with H&E (black arrow indicate lesion border) or toluidine blue (red arrows indicate mast cells), scale bars: upper panels, 500 µm; middle and lower panels, 50 µm; Morphometry of (c) leukocytes, (d) fibroblasts and (e) mast cells, 7 days after ear lesions; (f) Photomicrographs of the back skin, 7 days post-injury, stained with H&E (black arrows indicates lesion border) and Toluidine blue (green arrows indicate mast cells), scale bars: upper panels, 200 µm; middle and lower panels, 50 µm; Morphometry of (g) leukocytes, (h) fibroblasts and (i) mast cells, 7 days after lesions on the back skin. Data represent mean ± SEM, *P ≤ 0.05 compared with saline group (N=5).

### Intraperitoneal immunization and induction of systemic effects of oral tolerance

On the seventh day after oral treatment and 15 minutes before injuries, mice in the oral OVA-treated group (tolerant) and a control immune group received an i.p. injection of 10 μg of ovalbumin in 1.6 mg of Al(OH)_3_. A control group (adjuvant) received only the adjuvant Al(OH)_3_ and another group (saline) received saline.

### Injuries on the skin of the back and in the ears

Mice were anesthetized by i.p. injection of xylazine (16,5 mg/kg) and ketamine (97 mg/kg) diluted in saline and subjected to concomitant injuries on the skin of the back and in the ears. The skin on the dorsal region of the trunk was shaved and two excisional injuries were made using a 6mm diameter surgical punch. An injury was made in the center of each ear with a 4mm diameter surgical punch. Until recovery from anesthesia, the mice were kept blindfolded with cotton wool soaked in physiological saline to prevent corneal drying. After the injuries, the animals were kept in individual cages until the day of euthanasia.

### Preparation of samples for histological analysis

At 7 and 40 days after the injuries (Figures 1a and 2a), the animals were anesthetized with xylazine and ketamine and euthanized by cervical dislocation. Tissue samples containing the region of lesions were collected and fixed in Carlson formalin in Milloning buffer (pH 7.0) for 24 hours and then transferred to 70% alcohol for at least 1 hour before histological processing and inclusion in paraffin. The samples (N=5 per tissue, per time point) were sectioned at 5 μm thickness using a semi-automatic microtome (Microm HM315, GMI) and stained with Hematoxylin and Eosin (H&E), Toluidine Blue, Massom’s Trichrome, Pricrossirus red and Resorcin Fuchsin.

### Histological analysis

Histological sections were analyzed by two investigators, in a double-blind manner, using a light microscope (Olympus BX 40, Tokyo, Japan) or a polarized light microscope (Olympus, BX43, Tokyo, Japan). The cells were counted by two investigators, in a double-blind manner, using an intersection grid attached to the ocular lens (Thomas Scientific, Swedesboro, NJ). Leukocytes, mast cells and fibroblasts were identified by their characteristic color and morphology and counted in 10 fields of 10,000 μm2 each, within the area of the lesions and the results expressed as mean ± SEM.

### Score of cutaneous scar

After microscopic analysis of the dorsal skin stained with Masson’s Trichrome at 40 days post-injury, we determined a score to compare the scars of the different groups. The scars were categorized as severe, moderate or mild according to the intensity of deposition and organization of the collagen fibers. Scars were considered severe when the neodermis contained thinner, confluent collagen fibers parallel to the epidermis. In contrast, scars classified as mild showed thicker, intertwined and closely packed collagen fibers, similar to the pattern observed in intact skin, but without epidermal attachments, such as hair follicles. Moderate scars presented a mixture of thin and thick collagen fiber. The scars were scored by two independent investigators and the results expressed in values for each category (1-2 for mild, 3-4 for moderate and 5-6 for severe scar).

### Statistical analysis

Were performed using Graphpad Prism software (GraphPad Software, CA). The significance of differences between groups was determined using the One-Way ANOVA test followed by the Bonferroni test. Values of P ≤ 0.05 were considered significant.

## Results

### Intraperitoneal injection of ovalbumin (OVA) with Al(OH)_3_ adjuvant in mice that drank OVA reduced the inflammatory infiltrate in concomitant wounds in the ears and on the back skin

At 7 days after the injuries (Figure 1a), histological analyses of the wound beds in the ears of the animals that drank OVA showed a mild to moderate inflammatory infiltrate, with complete re-epithelialization and no cartilage at the injury site (Figure 1b). The saline, adjuvant and immune control groups presented intense inflammatory infiltrate and, in addition, presented hyperplastic epithelium, vascular congestion and edema. The control groups also did not present cartilage at the injury site. Corroborating the qualitative analyses, morphometric analyses regarding the number of inflammatory cells in the wound beds of the ears showed a lower number of leukocytes and mast cells in the group that drank OVA compared to the control groups (Figure 1c-e).

Regarding the wounds on the skin of the back, qualitative histological analyses at day 7 after the injuries showed that the group that drank OVA presented less inflammatory infiltrate, in addition to having complete re-epithelialization, less vascularization and less presence of adipocytes in the wound bed compared to the control groups (figure 1f). Morphometric analysis of the number of inflammatory cells and fibroblasts in the wounds on the skin of the back showed that the group that drank OVA had a lower number of leukocytes, mast cells and fibroblasts compared to the control groups (Figure 1g-i).

### Intraperitoneal injection of OVA plus Al(OH)_3_ adjuvant in mice that drank OVA improved wound healing in the ears and on the back skin

At 40 days after ear injuries (Figure 2a), a region of chondrocytes was observed in the animals that drank OVA that had formed beyond the site of the cartilage injury, as highlighted by the red dotted line in the samples stained with Masson’s Trichrome (Figure 2C). This characteristic was not observed in the animals in the saline group, but was observed to a lesser extent in the immune group. Furthermore, the group that drank OVA presented less evident fibrous scar tissue when compared to the control groups, in addition to the formation of chondrocyte aggregates in the region closest to the neoepithelium, visualized in Masson’s Trichrome and H&E staining.

**Figure 2.**
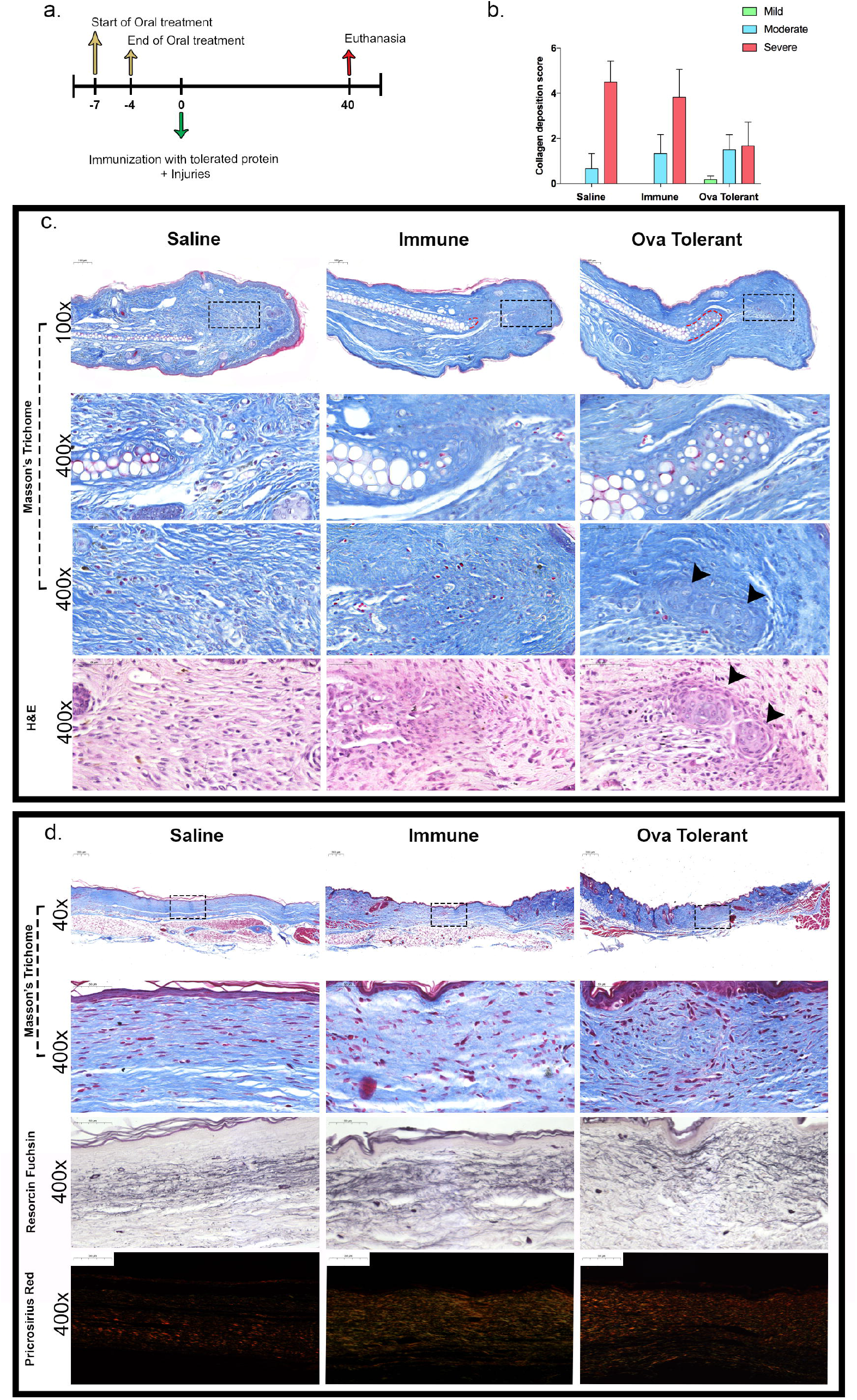
The i.p. injection of OVA in animals fed with this protein improved the healing of wounds in the ears and back skin. (a) Experimental design for analysis at day 40; (b) Collagen deposition score in samples analyzed 40 days after injuries on the skin of the back (N=5); (c) Photomicrographs of the ear with the scar, stained with Masson’s Trichrome or H&E. Red dotted line in low magnification Masson’s Trichrome staining indicate chondrocytes formed from the site of cartilage injury; boxes in low magnification Masson’s Trichrome staining indicate the area where the region are shown in high magnification to show chondrocyte aggregates in OVA tolerant mice; 400x magnification shows details of newly-formed chondrocytes extending from the original cartilage and chondrocyte aggregates (black arrows); (d) Photomicrographs of the back skin 40 days after injuries, stained with Masson’s Trichrome to evidence collagen fiber, Weigert’s Resorcin Fuchsin to show elastic fiber or Picrosirius Red to compare collagen fiber thickness.

At 40 days after the wounds on the skin of the back, histological analyses (figure 2d) of the scars of the animals that ingested OVA revealed deposition of thick, intertwined collagen fibers, with few non-fibrous components of the extracellular matrix, characteristics similar to those found in uninjured skin. On the other hand, in the scars of the saline and immune control groups, the collagen fibers were loosely organized, thinner and parallel to the epidermis, typical characteristics of scar tissue. Furthermore, analysis performed after staining with Picrosirius Red, which allows a qualitative analysis of collagen fibers so that thicker collagen fibers appear in redder tones when analyzed under a polarized light microscope, showed that animals that drank OVA had a greater quantity of reddish fibers, a characteristic similar to that of uninjured skin (figure 2d). On the other hand, control animals had a greater quantity of greenish fibers, a characteristic of thinner fibers normally found in scars. In addition, analysis of the elastic fibers 40 days after lesions on the skin of the back, using resorcin fuchsin staining (figure 2d), showed that in the animals that drank OVA, but not in the control groups, there was a better reorganization of the elastic fibers, so that they appeared homogeneously distributed throughout the neodermis, a pattern similar to the elastic fibers found in uninjured skin.

Comparisons of the scars on the skin of the back, using a score, showed that in the control groups the scars were predominantly severe, while in the group that drank OVA the scars were moderate or mild (Figure 2b).

## Discussion

Herein we showed that the inflammatory infiltrate triggered by multiple wounds, made simultaneously in different parts of the body, was reduced by the injection of a protein that had been previously ingested. Furthermore, there was better reorganization of the extracellular matrix in the scars of the animals that received the injection of the protein that had been previously ingested. The absorption of proteins by the gut mucosa normally leads to the establishment of oral tolerance, a specific immunological phenomenon that inhibits immune responses to the ingested proteins even when they are injected together with adjuvants ^18^. Previous work has shown that the injection of tolerated proteins does not result in immune responses but produces systemic and anti-inflammatory effects ^19^. Besides, other works have already shown that the systemic effects of oral tolerance, triggered by intraperitoneal or subcutaneous injection of dietary proteins, modify the initial phases of skin repair and reduce scars ^20,21^. This is the first work to show that the systemic effects of oral tolerance can improve the healing of concomitant wounds made in different parts of the body.

One of the characteristics of wound repair modified by the systemic effects of oral tolerance was the reduction in the number of mast cells in the lesion area, as evidenced in Figure 1. Mast cells can produce a large number of growth factors and cytokines and the action of these cells during wound repair may involve multiple mechanisms ^22^. Mast cells are involved in all phases of repair, from the initial inflammatory phase to collagen deposition by influencing the activity of fibroblasts and different studies have shown an increase in the number of mast cells in hypertrophic scars ^22^. Other studies have shown that a decrease in mast cells in the injured area correlated with better healing ^23^. During the inflammatory phase, factors produced by mast cells promote the recruitment of neutrophils and macrophages and blocking mast cell activation in the early phase of repair reduced skin scarring ^24^. Furthermore, mast cells may interact directly with fibroblasts, promoting their activation and the production and remodeling of the extracellular matrix ^25^.

Different studies have shown that reducing the inflammatory phase of repair correlates with reduced scarring and regeneration ^5^. Our results showed a reduction in the number of inflammatory cells at 7 days after the injuries and better reorganization of collagen fibers and elastic fibers at 40 days after the injuries in the skin of the back. Furthermore, in the injuries in the external ear (auricle) there was a reduction in the number of inflammatory cells that resulted in a less fibrotic repair, including the formation of chondrocyte aggregates. The presence of chondrocyte aggregates suggests that the systemic effects produced by the injection of tolerated proteins may benefit the regeneration of a tissue (the cartilage) that normally does not present a good regeneration potential. These results, together with previously published results showing a positive effect of the injection of tolerated proteins in the repair of bone defects ^17^, suggest a great potential for the use of this technique in the treatment of osteochondral defects. Further studies need to be conducted to translate the results obtained with animal models to the clinic.

Less fibrotic repair may result from the reduction of the inflammatory infiltrate itself, since the actions of inflammatory cells can produce lesions in the surrounding tissue ^26^. Furthermore, a greater inflammatory infiltrate may delay the resolution of inflammation and hinder the reorganization of damaged tissue ^27^. However, our results do not exclude the action of the systemic effects of oral tolerance, produced by the injection of tolerated proteins, on the migration and activation of fibroblasts.

Herein we did not study the specific mechanisms triggered by injection of the tolerated protein. However, previous works have shown that the systemic effects of oral tolerance, produced by injection of the ingested protein, produce changes in the expression of adhesion molecules and in the kinetics of cytokines, which may explain the reduction of the inflammatory infiltrate in the wound bed ^15,28^.

Previous studies have shown that i.p. injection of a protein previously given orally, concomitantly with intravenous injection of Schistosoma mansoni eggs, reduces the formation of pulmonary granulomas ^16^. A recently published study showed that i.p. injection of a protein present in the rat diet, concomitantly with a bone fracture, reduced the inflammatory infiltrate in the lesion and promoted better bone repair ^17^. The demonstration, through different studies, that the systemic effects of oral tolerance can reduce the inflammatory infiltrate in lesions of different organs and subsequently reduce fibrosis, suggests a significant clinical application. Several factors can generate multiple injuries in an individual, such as surgeries, traffic accidents, falls or sports injuries^1,29^. The limitations for the clinical application of the technique we present in our study is that it has not been tested in humans. However, the potential for medical application of dietary protein injection to improve multiple wound healing is very attractive and deserves to be further investigated.

## Conclusion

Oral administration of ovalbumin followed by the injection of this protein concomitantly with several wounds made in different body parts resulted in a reduction in the inflammatory infiltrate and improved the wound healing, favoring to the formation of less pronounced scars.

## Acknowledgments

We gratefully acknowledge funding from the Fundação de Amparo à Pesquisa de Minas Gerais – MG, Brasil (FAPEMIG, grant number APQ-00429-22) and Coordenação de Aperfeiçoamento de Pessoal de Nível Superior – Brasil (CAPES, Finance Code 001). I.B.C.N and A.V.S.A. received scholarchips from CAPES and FAPEMIG, respectively.

